# Resistome SNP Calling via Read Colored de Bruijn Graphs

**DOI:** 10.1101/156174

**Authors:** Bahar Alipanahi, Martin D. Muggli, Musa Jundi, Noelle Noyes, Christina Boucher

## Abstract

**Motivation:** The resistome, which refers to all of the antimicrobial resistance (AMR) genes in pathogenic and non-pathogenic bacteria, is frequently studied using shotgun metagenomic data [14, 47]. Unfortunately, few existing methods are able to identify single nucleotide polymorphisms (SNPs) within metagenomic data, and to the best of our knowledge, no methods exist to detect SNPs within AMR genes within the resistome. The ability to identify SNPs in AMR genes across the resistome would represent a significant advance in understanding the dissemination and evolution of AMR, as SNP identification would enable “fingerprinting” of the resistome, which could then be used to track AMR dynamics across various settings and/or time periods.

**Results:** We present LueVari, a reference-free SNP caller based on the read colored de Bruijn graph, an extension of the traditional de Bruijn graph that allows repeated regions longer than the *k*-mer length and shorter than the read length to be identified unambiguously. We demonstrate LueVari was the only method that had reliable sensitivity (between 73% and 98%) as the performance of competing methods varied widely. Furthermore, we show LueVari constructs sequences containing the variation which span 93% of the gene in datasets with lower coverage (15X), and 100% of the gene in datasets with higher coverage (30X).

**Availability:** Code and datasets are publicly available at https://github.com/baharpan/cosmo/tree/LueVari.

## 1 Introduction

Antimicrobial resistance (AMR) refers to the ability of an organism to persist in the face of exposure to an antimicrobial agent, i.e., an antibiotic. The *resistome*, which refers to the set of all AMR genes found in pathogenic and non-pathogenic bacteria, defines the potential resistance to known antibiotics. Shotgun metagenomic data has already been generated to characterize the resistome in clinical [14, 47] and food production [38] settings. The characterization of various resistomes includes identification of specific AMR genes and a measure of their abundance. One element of the resistome that has not yet been sufficiently characterized is the profile of single nucleotide polymorphisms (SNPs) within the AMR genes that comprise any given resistome. Characterization of the SNP profile would be very informative, as such a profile would allow specific AMR genes to be tracked across various settings and time points. Such *traceability* would greatly advance our understanding of how AMR disseminates and evolves; however, the ability to perform such traceability studies hinges on the accurate identification of AMR SNPs within metagenomic data. While methods do exist for identification of SNPs within eukaryote species, few current methods are suitable for metagenomic data—a sentiment expressed by Nijkamp et al. [37] when they state: “..there is a lack of algorithms for finding such variation in metagenomes.”

In this paper, we develop a scalable, reference-free method for identifying SNPs in metagenomic data, which we call LueVari. In order to be suitable for metagenomics, it is important that any SNP calling method be reference-free, as only a small fraction of the total diversity of microbes are culturable [44] and therefore, the majority of organisms within a given metagenomic sample will likely not have a reference genome in the near future. Hence, reference-guided tools will only detect a small fraction of available SNPs in metagenomes. Current reference-free methods specifically designed for metagenomic data typically require an “assembly” step that uses an overlap-layout-consensus (OLC) approach [18] or an Eulerian (de Bruijn graph) paradigm [3, 27, 34, 23, 45]. SNP detection in metagenomic data is challenging because it exacerbates the weaknesses of these two algorithmic approaches. De Bruijn graph-based SNP callers require each read to be fragmented into *k*-length sequences (called *k*-mers), and are thus prone to inappropriate combining of read segments from different species, resulting in *chimeric sequences*. OLC approaches–such as Bambus2 [18]–have an advantage over de Bruijn graph approaches in that they build an overlap graph in which the entirety of a read (rather than read *k*-mers) corresponds to a single node in the graph. As a result, sequences and SNPs recovered from these graphs are supported by collections of entire reads, rather than *k*-mers—an attribute of overlap graphs known as *read coherence* [33]. In theory, this reduces the frequency of chimeric SNPs. Unfortunately, OLC approaches suffer from computational inefficiency and thus are unable to handle large datasets such as metagenomic sequence datasets [25].

De Bruijn graph approaches have greater capacity to scale to larger datasets, especially in light of existing succinct data structures for representing de Bruijn graphs [43, 7, 8, 48, 4]. However, current de Bruijn graph-based methods lack the read coherence needed to detect SNPs accurately. Thus, there currently exists a significant trade-off between only being able to call SNPs within short sequences constructed from non-branching paths, or risk of chimeric sequences that arise from spanning branches in the graph. Furthermore, the potential for chimeric sequences becomes more frequent in resistome analysis due to sequence homology between AMR genes. Specifically, AMR genes within the same AMR class can share homologous regions that are typically between 60 bp and 150 bp in length [20], which is longer than the typical *k*-mer value and shorter than the read length. Hence, such homologous regions often correspond to several connected paths in the de Bruijn graph that are difficult to traverse unambiguously, implying that the de Bruijn graph cannot be used to reliably and correctly reconstruct the sequences corresponding to these AMR genes and their corresponding SNPs. As previously mentioned, these paths are more likely to be read coherent in the overlap graph, but the time complexity of OLC methods is too large to be practical for large-scale resistome analysis, which is likely to encompass the detection of AMR genes and their SNPs in hundreds or thousands of samples. For example, the USDA is calling for food production facilities to phase in use of sequencing to monitor AMR by 2021—if accomplished, even at a small scale, thousands of samples will need to be sequenced and analysed to monitor food-borne outbreaks.

Therefore, an approach for identification of SNPs in AMR genes within metagenomic data necessitates a method that combines the scalability of de Bruijn graph approaches with the read coherence of OLC approaches. To address this need, we develop a de Bruijn graph based SNP caller. It extends the concept of the colored de Bruijn graph, which was first introduced by Iqbal et al. [15] for the detection of variants in eukaryotic species. Given a set of *n* samples, the colored de Bruijn graph extends the traditional de Bruijn graph in that each node (and edge) in the graph has a set of associated colors in which each color corresponds to one of the *n* samples. In Iqbal et al.’s [15] original application, each sample corresponded to the sequence data of one individual and traversal of the colored de Bruijn graph allowed for sequence variation to be detected, along with the individuals containing that variation. Although the colored de Bruin graph allows for detection of genetic variation among individuals of a population, it lacks read coherence. In order to overcome this limitation, we propose an approach that we term a *read colored de Bruijn graph*. Briefly, a read colored de Bruijn graph annotates each node (and edge) by an unique color that corresponds to each individual sequence read (in one or more samples), allowing for read coherence to be preserved among paths longer than the *k*-mer size (typically, *k ≤* 60 bp). We formally define this concept later in this paper.

However, it does present construction challenges not present in colored de Bruijn graphs. One such challenge is that a metagenomic sample may be too large to store on even the largest servers’ hard drives in uncompressed form. For example, a set of metagenomic samples from a cattle production facility [38] contains close to 41 billion 32-mers, with the first sample containing over 57 million reads^1^. Storing each *k*-mer-read combination with a single bit would require 285 petabytes of space. This mandates that the succinct representation be built in an online fashion such that the complete uncompressed matrix need never be stored explicitly. Therefore, we present a succinct data structure to construct and store the read colored de Bruijn graph, which extends the representation of Muggli et al. [32]. *Our main contributions*. In this paper, we define the read colored de Bruijn graph along with several new concepts and demonstrate how it helps resolve chimeric sequences. This is allows LueVari to not only report the SNP but also correctly reconstruct the sequence containing the SNP. This is vital to metagenome applications where a reference genome is frequently unknown. Hence, ours was the only method to output sequences that span between 93% to 100% of the gene–even in the case of shallow coverage. For example, when the sequence data corresponds to 30X coverage and the SNP rate was 0.005, LueVari constructed all genes containing SNPs whereas DiscoSNP reported sequences that covered 71% (on average) of the gene, and Bubbleparse reported sequences that covered 41.76%. In addition, when compared to current state-of-the-art methods, LueVari was the only method that had reliably high sensitivity (between 73% and 98.5%); DiscoSNP and Bubbleparse had sensitivity between 32.8% and 91%, and 75% and 85%, respectively. Lastly, we show LueVari has the scalability to analyze all the data from a large-scale data collection effort aimed at characterizing the resistome of a commercial food production facility in the United States.

## 2 Related Work

As previously mentioned, there are both reference-based and reference-free SNP and variant callers. the majority of the SNP callers are developed for diploid organisms, namely eukaryotes, which limits their effectiveness on prokaryotes [51]. However, there do exist some methods that aim to detect variation in metagenomes. Among the reference-based variant callers, there are those that first align reads to the reference and then process this alignment. These methods include Hansel and Gretal [36], LENS [19], Platypus [41], MIDAS [35], Sigma [1], Strainer [11], and ConStrains [26]. Similar to these algorithms, are reference-based read alignment methods that use combinatorial optimization techniques to find sequences that best explain the reads and thus, can be used for variant detection in metagenomic samples. These methods include QuRe [40], ShoRAH [50], and Vispa [2]. The combinatorial optimization approaches are computationally intensive, limiting their applications to only relatively small datasets [13].

There are several reference-free variant callers. MaryGold [37], Bubbleparse [22], DiscoSNP [46],metafast [45], crAss [9], Commet [29], compareads [28], and FOCUS [42] are all reference-free. MaryGold [37], Bubbleparse [22] and DiscoSNP [46] are comparable to LueVari in that they are able to detect variants in metagenomes without a reference. There are several algorithms that do comparative metagenomics and return a similarity measure rather than specific variants that merit mention. These include metafast [45], crAss [9], Commet [29], compareads [28], and FOCUS [42]. Lastly, there are several reference-free variant callers that designed for a specific application and not directly comparable to LueVari, including KSNP3 [12], NIKS [17], Stacks [6] and 2k+2 [49]. Stacks [6] is designed for restriction enzyme based sequencing protocols. NIKS [17] is The read colored de Bruijn graph is an attractive concept because it avoids chimeric sequences by maintaining each read as a separate color. designed for whole-genome sequencing protocols. KSNP3 [12] and 2k+2 [49] are designed to detect the variations between datasets.

## 3 Definition of Read Coloring

As we previously discussed, de Bruijn graph approaches for SNP calling lose read information when the reads are fragmented into *k*-mers, which introduces the possibility of the sequences containg the SNP not being read coherent [33]. This lack of read coherence exacerbates the difficulty of detecting SNPs in metagenomes. In this section, we formally define the concept of the read colored de Bruijn graph and show how it is used by LueVari to detect SNPs accurately.

### 3.1 Read Colored De Bruijn Graphs

We begin by defining the (traditional) de Bruijn graph. We let *R* = {*r*_1_, …, *r_n_*} be the set of *n* input reads. We construct the de Bruijn graph for *R* by creating an edge for each *k*-mer in *R*, labelling the nodes of that edge as the (*k −* 1)-mer prefix and (*k −* 1)-mer suffix of that *k*-mer, and lastly, gluing nodes that have the same label. Here is an example of gluing: if node *v*_1_ with label ACG has outgoing edge with label C and node *v*_2_ with the same label ACG, has outgoing edge with label G, since both of nodes have same label, to make sure that all of labels in de Bruijn graph are distinctive, we glue them which means that the outgoing edge of *v*_2_ will be added to *v*_1_, hence *v*_1_ has two outgoing edges with labels C and G, and *v*_2_ will be deleted. The story is the same for incoming edges. Next, we define the concept of a *sub-read*, which is necessary for defining the read colored de Bruijn graph. Given a read *r* of length *l* it follows that there are *l – k* +1 *k*-mers.We denote the *k*-mers of *r* as 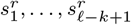 We define the *sub-reads* of *r* as the sets of *k*-mers, denoted as 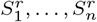, (*n* ≤ *l* + *k* + 1), where the following is true: (1) every *k*-mer of *r* is contained in one 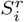, and (2) 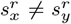 for all 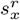 and 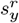 contained in the same set 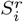.

More intuitively, we construct the set of sub-reads of *r* by creating *k*- mers from r until a repeated *k*-mer occurs; then, when we see a repeated *k*-mer, those *k*-mers are grouped into one sub-read and a new sub-read for r is created.We continue this process until all *k*-mers are added to a set.We note that if there are no repeated *k*-mers in *r* then the set of *k*-mers itself is the sub-read of *r*.We nowgive an an example to illustrate this: let *r* be equal to ACGTACGTACGT and *k* = 3. The substring ACGT is repeated three times in *r* and therefore, the sets of sub-reads are 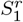 = {ACG, CGT, GTA, TAC}, 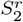 = {ACG, CGT, GTA, TAC}, and 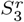 = {ACG, CGT}. We note that making sub-reads in the case of repeated *k*-mers will disambiguate the traversal of whirls (directed cycles) that have length greater than or equal to *k* and less than or equal to *l*. For instance in above example, the read ACGTACGTACGT, will make a whirl at node ACG that makes the traversing ambiguous. Note that this problem will be solved if we follow the sub-reads 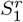 and 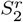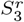 which lead us to traverse ACG three times and finish the traversing at CGT.

Next, we define the read colored de Bruijn graph for a set of reads *R* constructively as follows: (1) we create the sub-reads of *r* for all *r* ∊ *R* (denote d as 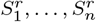,*n* ≤ *l* – k + 1.) (2) we assign a color 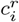 for every sub-read 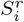, and (3) we build the de Bruijn graph as above with the modification that color 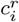 is associated every edge e if the *k*-mer corresponding to *e* is contained in 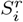 Hence, we view the read colored de Bruijn graph as a graph *G* = (*V;E*) and a binary matrix *C*, where there exists a row for each distinct *k*-mer, and a column for each sub-read and *C*(*i; j*) = 1 if the *k*-mer associated with edge *e_i_* ∊ *E* is present in *j*th sub-read; and *C*(*i; j*) = 0 otherwise. We refer to *C* as the color matrix. We illustrate a read colored de Bruijn graph in Figure A.1 in the Supplement. Briefly, we mention that the addition of read coloring to the de Bruijn graph resolves repeats that are shorter than or equal to the read length, and longer than or equal to the *k*-mer length.We discuss this more in-depth in Subsection 4.1.

### 3.2 Multi-Colored Bulges

We define a *bulge* in *G* as a set of disjoint paths (*p*_1_, …, *p_n_*) which share a source and sink node, where we refer to paths (*p*_1_, …, *p_n_*) as the branches. Next, we define a path *p* = *e*_1_ *… e_f_* (*e_i_*s (1 *≤ i ≤ R*) are edges) in the read colored de Bruijn graph as *color coherent* if the sets of colors corresponding to *e_i_*, *e_i_*_+1_, *S_i_* and *S_i_*_+1_, are such that *S_i_ ∩ S_i_*_+1_ *I* = *∅*, for all *i* = 1, …, *R−*1. Thus, we refer to *multi-colored bulge* as a bulge where the branches are color coherent and also have disjoint lists of colors. We illustrate a multi-colored bulge in Figure 1. Lastly, we define an *embedded multi-colored bulge* in *G* as a multi-colored bulge that occurs in a branch of another multi-colored bulge.

**Fig 1.**
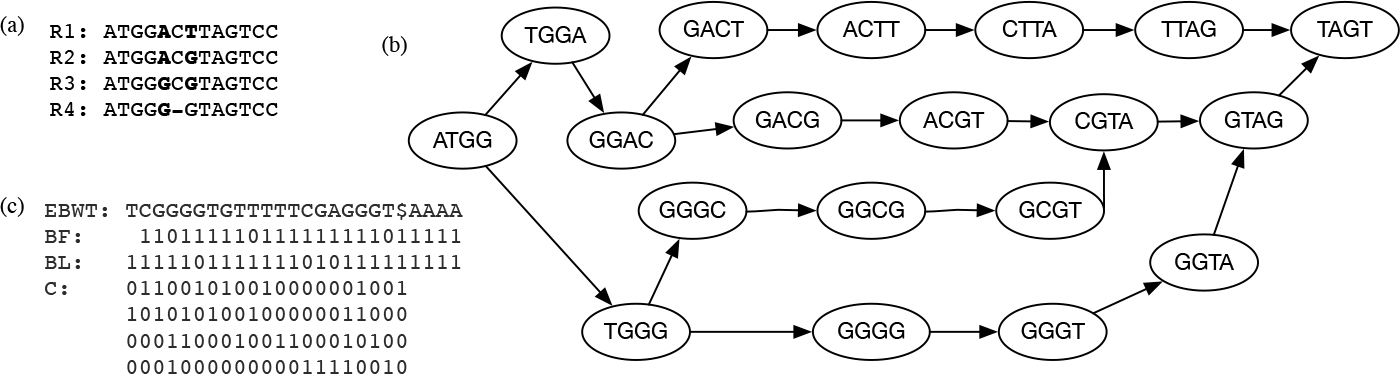
In this example, we illustrate a complex multi-colored bulge that arises from the existence of multiple SNPs in a single gene. In (a), we show the reads with multiple variations. In (b), we illustrate the de Bruijn graph which was constructed for the set of reads in (b). Lastly, in (c) we give the succinct representation of read-colored de Bruijn graph.

## 4 Methods

In this section, we go over the main steps of LueVari in detail.

### 4.1 Construction of the Read Colored de Bruijn Graph

We build the read colored de Bruijn graph for reads *R* by first constructing the de Bruijn graph for *R*, computing the sub-reads of *R*, and creating the color matrix (in a compressed format). We use the BOSS data structure [4], which is based on the Burrows-Wheeler Transform (BWT) [5] for constructing and storing the graph. Here, we will give a brief overview of this representation and we refer the reader to the original paper by Bowe et al. [4] for a more thorough explanation of this data structure

Our first step in constructing this graph *G* for a given set of *k*-mers is to add dummy *k*-mers (edges) which ensure that there exists an edge (*k*-mer) starting with first *k −* 1 symbols of another edges last *k −* 1 symbols and thus, that the label of each edge and node *G* can be recovered. After this small perturbation of the data, we construct a list of all edges sorted into right-to-left lexicographic order of their last *k −* 1 symbols (with ties broken by the first character). We denote this list as *F*, and refer to its ordering as co-lexicographic (colex order). Next, we define *L* to be the list of edges sorted co-lexicographically by their starting nodes with ties broken co-lexicographically by their ending nodes. Thus, we note that two edges with same label have the same relative order in both lists; otherwise, their relative order in *F* is the same as the lexicographic order of their labels. The sequence of edge labels sorted by their order in list *L* is called the *edge-BWT* (EBWT). Now, we let *B_F_* be a bit vector in which every 1 indicates the last incoming edge of each node in *L*, and let *B_L_* be another bit vector with every 1 showing the position of the last outgoing edge of each node in *L*. Given a character *c* and a node *v* with co-lexicographic rank rank(*c*), we can determine the set of outgoing edges of *v*, using *B_L_* and then search the EBWT(*G*) for the position of edge *e* with label *c*.We can find the co-lexicographical rank of outgoing edge of *e* using *B_F_*, and thus, traverse the graph by repeating this process,.

After we construct the BOSS representation of the de Bruijn graph on *R*, we align all *k*-mers to the union of all the sub-reads using Bowtie [21] in order to find all the sub-reads that contain each of *k*-mers. (We only consider perfect matches). Then we store the *k*-mers and their lexicographical order in a map *M* so it can be used for the construction of the color matrix. We use Elias-Fano vector encoding [10, 30, 39] to store *C* since it permits on-line construction as long as all “1” bits are added in increasing order of their index in the vector. For example, we cannot fill column six of *C* before column five is filled. Therefore, we build the color matrix by initializing each position to “0” and then updating each row at a time. We recall that each row corresponds to a *k*-mer and the rows are sorted in lexicographical order. Thus, we first find its lexicographical ordering of a *k*-mer *s_k_* using *M*, which we denote as *i*, in order to find the row in *C* corresponding to *s_k_*. Using the alignment of *k*-mers to sub-reads, we store the indices of the sub-reads that contain *sk* in a vector 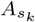. We sort 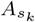and update the *i*th row of *C* in this order, i.e., *C*(*i, j*) = 1 for all *j*th sub-reads containing the *i*th *s_k_*. We store and sort the indices in this manner to ensure that we meet the construction requirement of Elias-Fano. After we construct 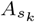in a compressed format, we append it to growing color matrix *C*. We continue with this process until all *k*-mers have been explored.

### 4.2 Search for Multi-Colored Bulges

Next, we traverse *G* in a read-coherent manner in order to determine all multi-colored bulges. In the first step of this traversal we iterate through all nodes in *G* and determine those that are potential source nodes of a multi-colored bulges, meaning that the out-degree is greater than one and the sets of colors of the outgoing edges are disjoint. Thus, given a node *v* with index *i* (in colex order), we determine whether *v* is a source node as follows. We first calculate out-degree of *v* using *B_L_* by finding *i*-th ‘1’ bit in *B_L_*, and counting the number of preceding ‘0’ bits *B_L_*. If the number of ‘0’ bits is *R* then the out-degree of *v* is *R* + 1. If the out-degree of *v* is not greater than one then we do not consider it further; otherwise, we determine the colors of its outgoing edges. We recall that the lexicographical order of the edges (*k*-mers) corresponds to their row index in the color matrix *C*. Therefore, if *v* has outgoing edges *e*_1_ and *e*_2_ with lexicographical order of *x* and *y*, respectively then we can determine the colors of *e*_1_ by finding all positions *R* where *C*(*x, R*) = 1 and those for *e*_2_ by finding all positions *R^I^* where *C*(*y, R^I^*) = 1. If these two sets are disjoint then *v* is a potential source node.

Next, we perform (modified) depth first search at each potential source node *v* to find one or more potential sink nodes. We modify the standard depth first search algorithm by adding a constraint to ensure all paths are color coherent, and by storing multi-colored bulges during the traversal. We ensure the paths of the search are color coherent by determining the intersection of the set of colors of the next outgoing edge with those of the previous one at each iteration, and terminating if there this intersection is empty. Further, we determine the in-degree of *v* at each iteration^2^, and store *v* as a potential sink node (along with the source node and branches) if the degree is greater than one. We note that we do not stop the traversal after encountering a potential sink node but instead, continue until all nodes reachable from *v* have been visited.

If we encounter a potential source node *u* while traversing a path *pv* starting at *v* then we determine whether *u* has been previously visited. If it has, then we search for *u* in the set of multi-colored bulges. If *u* is a source node of a multi-colored bulge with branches {*p_u_*_1_,…,*p_uf_*} and sink node *t_u_*, then we do not traverse *p_u_*_1_,…,*p_uf_*. Instead, we add each path in {*p_u_*_1_,…,*p_uf_*} to *p_v_* and continue traversing from *t_u_*. We note that *v* will have *R* branches {*p_v_* +*p_u_*_1_,…, *p_v_* +*p_uf_*} after concatenating the paths. If *u* has not been previously visited then depth first search is performed, starting with *u* as a potential source node, to determine whether there exists a multi-colored bulge with source node *u*. If there does not exist such a bulge then we resume the prior depth first search where *v* was the starting node. Otherwise, we store the multi-colored bulge (with source node *u*, set of branches {*p_u_*_1_,…, *p_uf_*}, and sink node *t_u_*), process the bulge in an analogous manner as just described, and resume the depth first search at *tu*. This ensures that we will detect all complex multi-colored bulges.

We illustrate this traversal using an example of a complex multi-colored bulge in Figure 1. We note that dummy incoming edges and one dummy outgoing edge (labeled with $) have been added to construct the succinct de Bruijn graph but since they do not clarify the traversal, we omit them in the depicted graph. In this example, we iterate through all nodes until we reach GGAC, which is the 6th node in colex order. In *B_L_*, the 6th ‘1’ bit has one preceding ‘0’ bit which means that GGAC has out-degree 2 since each ‘1’ bit in *B_L_* indicates the last outgoing edge. Now we determine if GGAC is a potential source node by first checking if its outgoing edges have disjoint color sets. We find the outgoing edges, which are *G* and *T*, by finding the range of EBWT[5, 6]. We determine the lexicographical order of GGACG which is 10. By considering the 10th column in color matrix *C^T^*, we can identify the sub-read(s) that GGACG comes from, which is only sub-read 2. By doing the same with the other edge GGACT of lexicographical order 11, we identify its supporting sub-read(s), which is sub-read 1. We witness that the color sets of outgoing edges are disjoint which implies GGAC is a potential source node. Hence, we start traversing at this node and follow the edge with label *G* (GGACG). For finding the destination node of this edge, we count the preceding edges in colex order. We witness that there are five edges with label *A*, two edges with label *C* in whole graph and we are at 4th *G* in EBWT. Thus, we count 1’s in *B_F_* [0, …, 11] and determine there are 10. This means that GGACG arrives at the 10th nodes in colex order that has incoming edges. Since first node ($$$$) has no incoming edge, GGACG arrives at 11th node in colex order. The node with colex order 11 is GACG. After finding the destination, we check the read coherency by finding the following edge, which is GACGT. As mentioned before, we can find the color set for this edge, which only includes sub-read 2. Since color sets of two consecutive edges have at least one intersection, traversing continues. Along the way nodes with in-degree greater than one will be saved as potential sink nodes (similar to what mentioned on *B_L_* to find out-degree, in-degree can be found using *B_F_*). In this example, TAGT is such a node. While traversing the other branch, if this node is visited again, the bulge will be detected and the start node, branches, and the end node will be stored. The exact same process will be performed on the second bulge with start node TGGG. Finally when the algorithm hits node ATGG, by hitting the already visited start nodes, it will not traverse the already known multi-colored bulges on nodes GGAC and TGGG again. By calling them from the list of multi-colored bulges, their branches will be added to the currently growing branches and by jumping to their end nodes, traversing finishes faster. Essential to our method is that, even though the multi-colored bulge at node ATGG comes from a variation of *A* and *G*, the reported branches, covers all embedded SNPs along the way, and for the complex bulge at start node ATGG, the total of four branches will be reported, which retrieve all four sub-reads with their SNPs.

### 4.3 Recovery of SNPs

Next, we process each multi-colored bulge *b* with branches {*p*_1_, …, *p_n_*} by recovering the longest color coherent path that occurs prior to the source node *s* by starting at *s* and travelling backward on the incoming nodes as long as their exists an unambiguous incoming edge, implying there exists one incoming edge (possibly part of a branch of an embedded multi-colored bulge) can be added to the current path and have it remain color coherent. If there exists such an edge then it is added to the current path and the traversal backward continues; otherwise the traversal is halted and the current path *ps* is saved. Similarly, a color coherent outgoing path is obtained from traversing the graph in a forward direction from the sink node *t*. We refer to this resulting path as *pt*. Lastly, *ps* is concatenated to each branch in {*p*_1_, …, *p_n_*}, *p_t_* is concatenated to each of these resulting paths, and their corresponding sequences are emitted. The SNPs in the sequences are recovered by alignment. This process is continued for all multi-colored bulges.

## 5 Results and Discussion

In this section, we compare the performance of LueVari to other SNP detection methods. In particular, we present the efficiency, sensitivity and precision of LueVari and the competing methods on simulated data, and demonstrate the scalability of LueVari by using it to identify distinct (“fingerprinted”) AMR genes in 34 samples taken from a food production facility that had previously-identified AMR genes. All experiments were performed on a 2 Intel(R) Xeon(R) CPU E5-2650 v2 2.60 GHz server with 1 TB of RAM, and both resident set size and user process time were reported by the operating system.

### 5.1 Results on Simulated Data

We note that majority of SNP calling methods output SNPs along with sequences flanking or containing the variant—the lengths of these sequences has an important role in SNP detection in metagenomes. The sequence must be long enough to locate the SNPs (e.g. gene and loci) in a unambiguous manner. This challenge is compounded in metagenomics variant detection, where the majority of bacteria is unculturable and unlikely to contain a reference genome. Therefore, we use the simulated data to evaluate the accuracy of LueVari and competing methods, as well as, to compare the relative length of the sequences outputted by the methods.

#### 5.1.1 Accuracy of SNP detection

We simulated four metagenomics datasets using BEAR, a metagenomics read simulator [16], in a manner that imitates the characteristics (number of reads, number of distinct AMR genes, and their copy number) of real shotgun metagenomics data generated for resistome analysis–namely those of Noyes et al. [38] and Gibson et al. [14]. We varied the copy number and number of paired-end reads for each dataset. Hence, in order to simulate a dataset with copy number *x* and *y* number of paired-end reads, we performed the following steps: (1) we selected 400 AMR genes from the MEGARes data at random without replacement, (2) we made *x* copies of each gene, (3) we added two copies of the *E. coli* K-12 MG 1655 reference genome, and two copies of the salmonella enterica subspecies I, serovar Typhimurium (S. typhimurium) reference genome, and lastly, (4) we simulated *y* paired-end reads from this resulting set of sequences. We used a 1% error rate for both these simulations. We note that all paired-end datasets contained 150 bp reads and 200 bp insert size. For this experiment, we simulated 270,598, 527,913, 1,110,150 and 2,110,753 paired-end short reads with 3, 5, 9, and 16 average copy number of the AMR genes, respectively. In brief, we call these datasets 270K, 500K, 1M and 2M. Please see the Supplement (Table A.7) for the results of comparison of sensitivity and precision of the reference free tools on datasets with same number of reads and varying number of gene copies.

We compared LueVari against DiscoSNP (DiscoSNP++) [46] and Bubbleparse [22], and note that the most recent version of these was used. Although MaryGold [37] and Bambus2 [18] are comparable methods, they are not available for current sequencing technologies. We also show the comparison of LueVari to SAMtools [24] and GATK [31] in the supplement (see Table A.4 and Table A.5) but note that these are reference-based methods. We used the MEGARes database as the reference. For example, we witnessed that GATK filtered 51.34%, 57.35%, 64.91% and 71.01% of reads in the 270K, 500K, 1M and 2M datasets, respectively, and thus, had low sensitivity for these datasets.

We summarize the results of this experiment for the reference-free methods in Table 1 and Table 2. We calculated the sensitivity and precision, based on the alignment of outputted sequences to the MEGARes database. LueVari and DiscoSNP are able to remove *k*-mers that have low multiplicity prior to SNP calling. Therefore, we ran DiscoSNP with two different thresholds: with the default setting (which is 3), and with the setting that achieve the highest performance (which is 0). For comparison, we ran LueVari with identical settings. We can see, with a slight penalty on precision and time, LueVari(0) has the highest sensitivity. Further, filtering the low abundant *k*-mers increased the precision of both LueVari and DiscoSNP; however, the sensitivity of DiscoSNP dropped remarkably (e.g. from 90.1% to 57.5%) whereas LueVari retained a high sensitivity (greater than 90% for all samples containing more than 500K reads). LueVari(3) had the highest sensitivity for three of the four datasets–with the one exception being the 270K dataset in which the sensitivity of LueVari is 2˜% less than Bubbleparse but has a precision that is 9.8% higher than Bubbleparse. We report the memory and time usage (CPU time) of all the methods in Table 2. All methods required less than 40 minutes and 12 GB of RAM on these datasets. DiscoSNP was the most efficient, yet the sensitivity was significantly lower than LueVari.

**Table 1.** In this table, we report the sensitivity and precision (reported as percentage) of all methods on the simulated datasets. We note the 270K, 500K, 1M and 2M datasets have 270,598, 527,913, 1,110,150 and 2,110,753 paired-end simulated reads, respectively. *k*=32 for all experiments except for Bubbleparse in which *k*=31 (*k* can not be even). We report in brackets the the sensitivity when interest is restricted sequences that have length greater than 200 bp.

|  | 270K |  | 500K |  | 1M |  | 2M |  |
| --- | --- | --- | --- | --- | --- | --- | --- | --- |
|  | Sensitivity | Precision | Sensitivity | Precision | Sensitivity | Precision | Sensitivity | Precision |
| LueVari(0) | 95.6 (**95.6**) | 11.2 | 96.2 (**96.2**) | 12.5 | 98.5 (**98.5**) | 5.6 | 97.5 (**97.5**) | 6.5 |
| DiscoSNP(0) | 81.7 (**59.1**) | 14.1 | 91.2 (**63.7**) | 14.2 | 90.1 (**56.1**) | 11 | 86.8 (**72.5**) | 12.5 |
| LueVari(3) | 73 (**73**) | 17.9 | 90.6 (**90.6**) | 16 | 94.6 (**94.6**) | 7.2 | 95 (**95**) | 6.3 |
| DiscoSNP(3) | 32.8 (**0**) | 26.7 | 67.5 (**0**) | 8.4 | 57.5 (**0**) | 15.0 | 55.6 (**0**) | 15 |
| Bubbleparse | 75.18 (**43**) | 8.1 | 85 (**67**) | 6.5 | 85.6 (**46.2**) | 4.4 | 85.6 (**49.4**) | 5.1 |

**Table 2.**
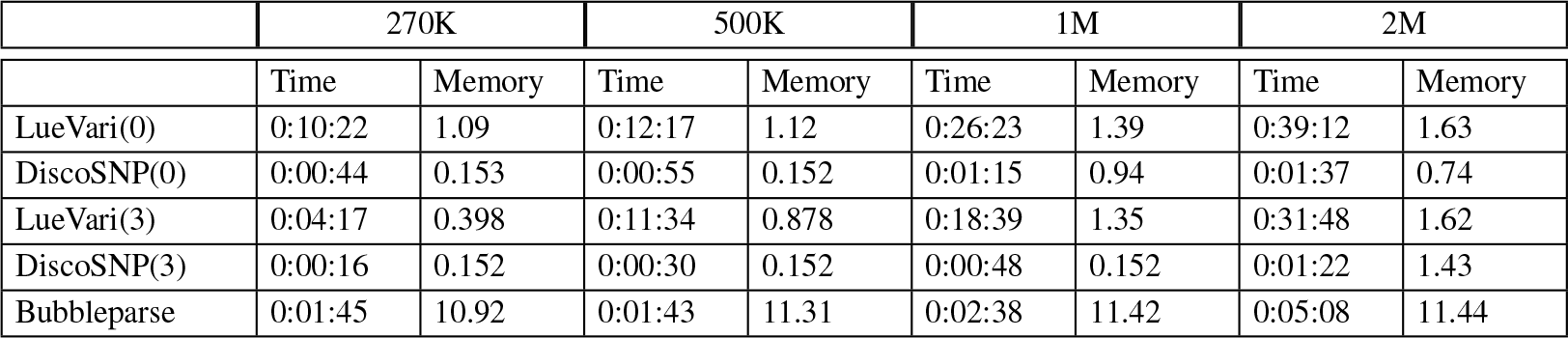
In this table, we give the performance results on the simulated datasets. 270K, 500K, 1M and 2M with 270,598, 527,913, 1,110,150 and 2,110,753 paired-end simulated reads. We report the peak memory in gigabytes (GB), and the running time as hh:mm:ss. *k*=32 for all experiments except for Bubbleparse in which *k*=31 (*k* can not be even).

|  | 270K |  | 500K |  | 1M |  | 2M |  |
| --- | --- | --- | --- | --- | --- | --- | --- | --- |
|  | Time | Memory | Time | Memory | Time | Memory | Time | Memory |
| LueVari(0) | 0:10:22 | 1.09 | 0:12:17 | 1.12 | 0:26:23 | 1.39 | 0:39:12 | 1.63 |
| DiscoSNP(0) | 0:00:44 | 0.153 | 0:00:55 | 0.152 | 0:01:15 | 0.94 | 0:01:37 | 0.74 |
| LueVari(3) | 0:04:17 | 0.398 | 0:11:34 | 0.878 | 0:18:39 | 1.35 | 0:31:48 | 1.62 |
| DiscoSNP(3) | 0:00:16 | 0.152 | 0:00:30 | 0.152 | 0:00:48 | 0.152 | 0:01:22 | 1.43 |
| Bubbleparse | 0:01:45 | 10.92 | 0:01:43 | 11.31 | 0:02:38 | 11.42 | 0:05:08 | 11.44 |

One important feature of LueVari is the ability to correctly reconstruct the sequences containing the SNPs. As shown in Table 1, when we only considered sequences that have at least 200 bp, we see that the sensitivity of the competing methods dropped dramatically. LueVari the sensitively of LueVari remained unchanged. This is an important feature since in metagenome application, a reference genome is frequently unknown. In this application, longer sequences are needed in order to unambiguously compare the SNP profiles between samples.

#### 5.1.2 Comparison of sequence lengths

We evaluate the ability of LueVari, DiscoSNP (DiscoSNP++) [46], and Bubbleparse [22] to detect SNPs in a unambiguous manner. To perform this comparison, we simulated four datasets using BEAR in an identical manner as described above, varying the SNP rate and copy number. First, we constructed two datasets with 258,180 and 2,991,107 paired-end sequence reads with mean copy number of 15 (range [10, 20]) and mean copy number of 30 (range [25, 35]), respectively. Here, we kept the SNP rate constant at 0.00125, and selected 16 AMR genes to be in the set. In addition, we simulated reads from *E coli* and salmonella in the manner that was described above. For simplicity we call these samples low-0.00125 and high-0.00125. Next, we simulated two additional datasets with 245,979 and 2,844,899 paired-end reads with mean coverage 15 (range [10, 20]) and 30 (range [25, 35]), respectively. Here, we set the SNP rate to 0.005, selected 10 AMR genes, and as in the previous experiment, included reads simulated from *E coli* and salmonella. We refer to these datasets as low- 0.005 and high-0.005.

By default DiscoSNP output sequences of length 2*k −* 1 where *k* is the value used to construct the de Bruijn graph, and the recommended length of sequences for Bubbleparse is 200 bp. Thus, we changed the parameters of DiscoSNP and Bubbleparse to output the longest possible sequences containing the identified SNPs. We ran DiscoSNP with -b2, and -T to ensure that it does not filter branching bubbles (bulges), and extends bubbles to unitigs/contigs. We ran Bubbleparse with -w 2,4000 to extract sequences around 4,000 nodes.

We report the results in Table 3 and refer the reader to the supplement for additional details about the results (Figure A.3 and Figure A.4). We define gene fraction to be the percentage of base pairs in a gene that are covered by the outputted sequence containing the identified SNP. Thus, we report the *mean gene fraction* (denoted as GF), which is mean of the gene fraction of all sequences (containing an identified SNP) outputted by a method. In addition, we report the percentage of genes that were *correctly recreated* (denoted as CRG), where we define a correctly recreated gene as one where the corresponding outputted sequence has 100% GF. If there are multiple unique SNP profiles for a single gene then this metric counts each profile as an individual gene that should be reconstructed. For example, if we have a dataset with a gene that has 8 different SNP profiles then there should be 8 genes outputted with 100% gene fraction—one for each unique SNP profile.

**Table 3.**
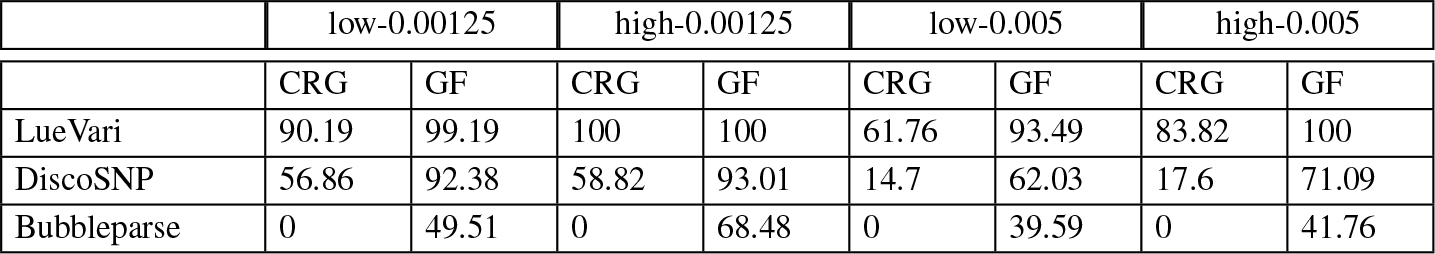
We give the metrics that describe the length of the sequences outputted by LueVari and competing methods. The low-0.00125 and high-0.00125 datasets contain a total number of 16 AMR genes with 0.00125 SNP rate. The low-0.05 and high-0.05 datasets contain a total number of 10 AMR genes with 0.05 SNP rate. *k*=32 for all experiments except for Bubbleparse in which *k*=31 (*k* can not be even).

We see that LueVari identifies the largest mean gene fraction. Hence, Bubbleparse, DiscoSNP and LueVari had 39.6% and 68.5%, 62% and 93%, and 93.5% and 100% as minimum and maximum GF, respectively. Furthermore, Bubbleparse, DiscoSNP and LueVari had a CRG as 0%, between 14.7% and 58.82%, and between 61.76% and 100%, respectively. These results reflect the algorithmic difference between the methods. In case of embedded bulges (where there exists a bulge(s) within bulge), LueVari reports all possible branching paths within a bulge, leading to a higher CRG. Whereas, competing methods only consider a single path within a bulge. This reflects the results in Table 3; the performance of DiscoSNP and Bubbleparse was degraded with a higher SNP rate which causes greater branching within the de Bruijn graph. LueVari was the only method that consistently had high mean gene fraction (greater than 93%), regardless of the coverage and SNP rate. This is an important feature since the resistome (and microbiome) consume a small amount of the biological sample and thus, will have very low coverage and varied polymorphism rate [38, 14]. Lastly, we note that although the gene fraction of DiscoSNP was relatively high for two of the four datasets we considered (e.g. 92% and 93% for low-0.00125 and high-0.00125, respectively) the CRG was low for all samples, ranging from 15% to 59%.

### 5.2 Results on Metagenomic Data from Food Production

We demonstrate the ability of LueVari to analyze real shotgun metagenomic data that were sequenced on an Illumina HiSeq 2500 system. The samples were selected across a beef production system, which contain different interventions (such as, high-heat and lactic acid treatment) aimed at decreasing AMR in consumable beef. Hence, this dataset is used to explore how microbial communities surrounding beef production facilities evolve in the presence of different food production interventions that aim to reduce pathogen load [38]. We ran LueVari on method on the shotgun metagenomic datasets, first filtering for eukaryote species (hen bovine and human DNA) and then filtering for *k*-mers that have low multiplicty, We report the number of the reads, distinct *k*-mers, detected bulges, total time of the pipeline, time for constructing the read-color matrix, time for the traversing, peak memory and size of the read-color matrix in Table A.6. As we can see in Table A.6, the parameters affect the traversal time are color matrix size, number of distinct *k*-mers, and number of bulges. For example, the largest traversing time belongs to sample 3, with largest number of bulges (23,782), largest color matrix (5.2 GB) and second greatest number of distinct *k*-mers (40,759,656). The size of color matrix depends on number of reads, number of distinct *k*-mers (size of de Bruijn graph) and total number of *k*-mers, which indicates the sparseness of the matrix (this value is not reported in the table). We emphasize that both the number of reads and number of *k*-mers of sample 1 are greater than those of sample 3, and still the color matrix of sample 3 is larger, which is due to a greater total number of *k*-mers on sample 3. In other words, the color matrix of sample 3 is more condensed. As one can see in Table A.6, with less than 21 GB of memory, and 6 GB of disk space, LueVari can scale for large datasets (more than 55 million read) in less than 19 hours.

## A.1 Supplement

### A.1.1 Illustration of the Read Colored De Bruijn Graph

In Figure A.1 we illustrate concepts related to the read colored de Bruijn graph, and their role in resolving cycles.

**Fig A.1.**
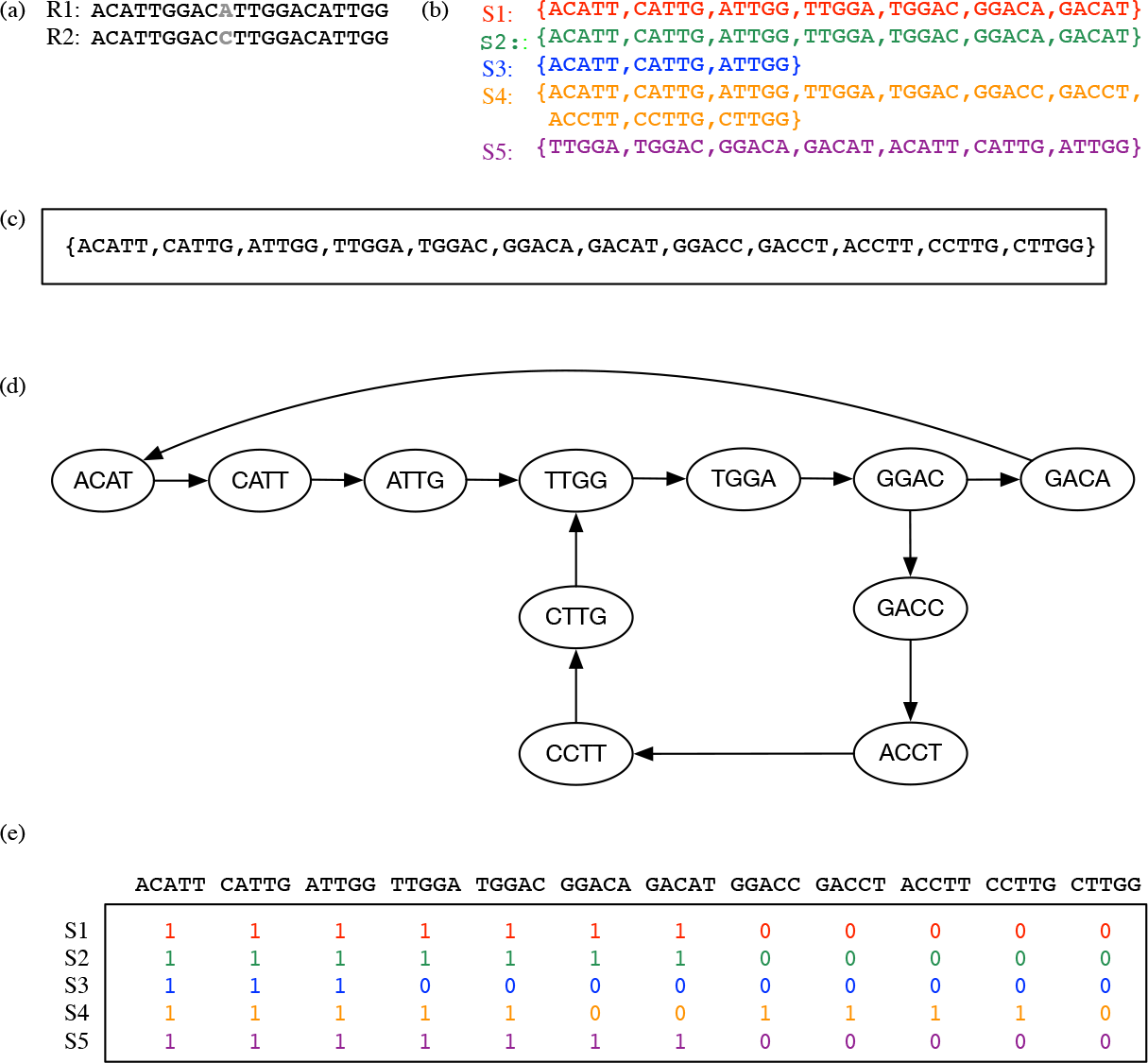
In(a) we illustration of two reads R1 and R2 representing a SNP (shown in grey). In (b), we show the sub-reads of R1 and R2; S1, S2, and S3 originate from R1, and S4 and S5 originate from R2. In (c), we give the *k*-mers that were constructed from R1 and R2. In (d), we show the de Bruijn graph constructed from the set of *k*-mers. And lastly, in (e), we show the color matrix constructed from the sub-reads and de Bruijn graph. If we traverse the read colored de Bruijn graph in a color coherent manner then ACATTGGACATTGGACATTGG and ACATTGGACCTTGGACATTGG are recovered. We note that we show the transpose of color matrix to save space.

### A.1.2 Gene Fraction Histogram for LueVari

In Figure A.2 we give a histogram of the gene fraction of the sequences outputted by LueVari.

**Fig A.2.**
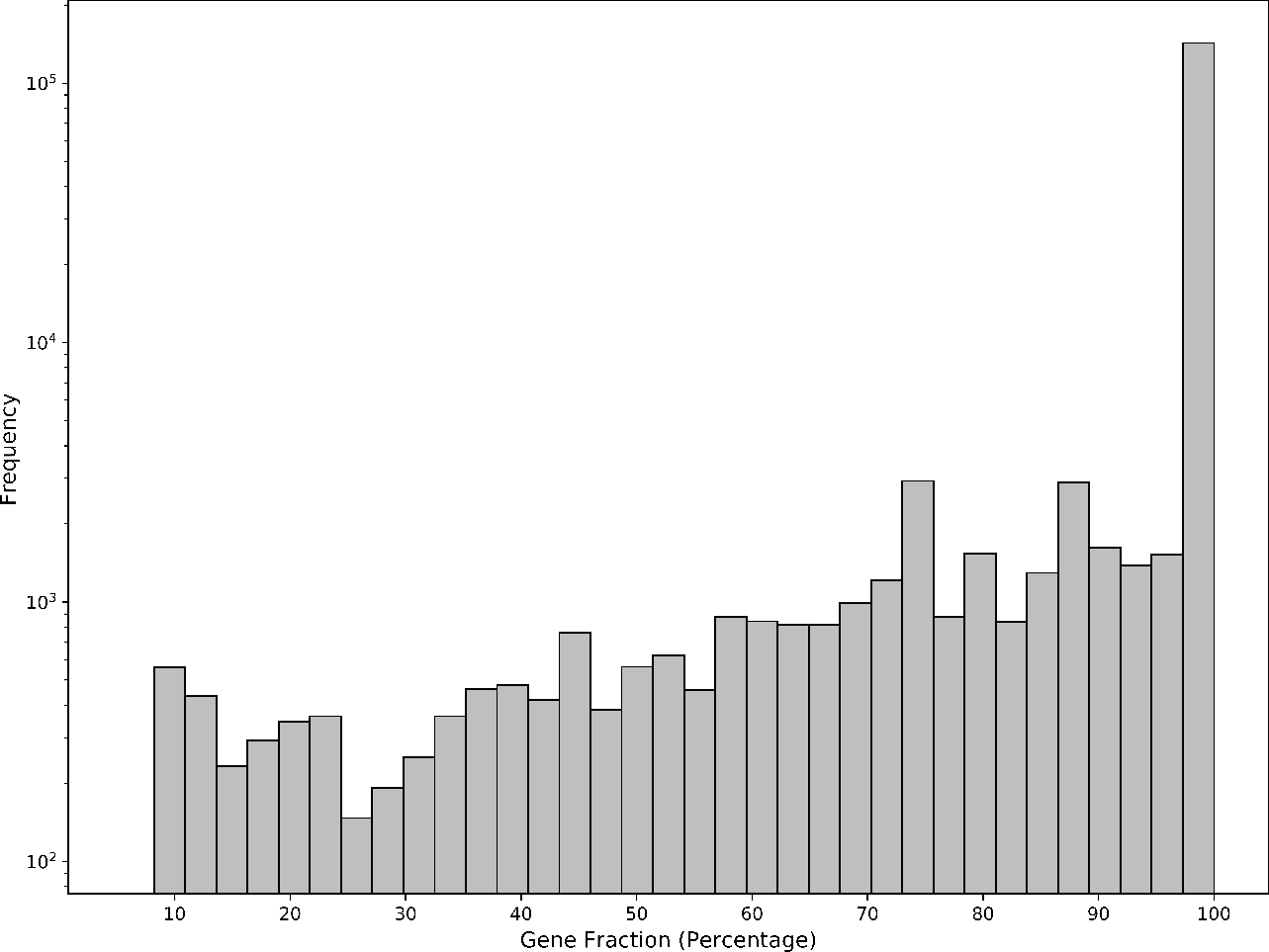
In this figure, we give a histogram of the gene fraction of the sequences outputted by LueVari. The y-axis gives the frequency in log scale. The x-axis gives the percentage of the gene that was covered by the sequence containing the detected SNPs. We note that this illustrates one of the main benefits of LueVari, which is that it allows the location of SNPs to be identified without disambiguity.

### A.1.3 Comparison of Gene fractions

We compared LueVari gene fraction with DiscoSNP and Bubblepars–two other reference free SNP callers, in four experiments. low-0.00125 and high- 0.00125 refer to simulated metagenomic read with SNP rate of 0.00125 and mean coverage of 15 and 30 respectively. Accordingly low-0.005 and high-0.005 refer to simulated metagenomic read with SNP rate of 0.005 and mean coverage of 15 and 30 respectively. As one can see mean gene fraction of LueVari is the highest among all tools. See graphs A.3 and A.4.

**Fig A.3.**
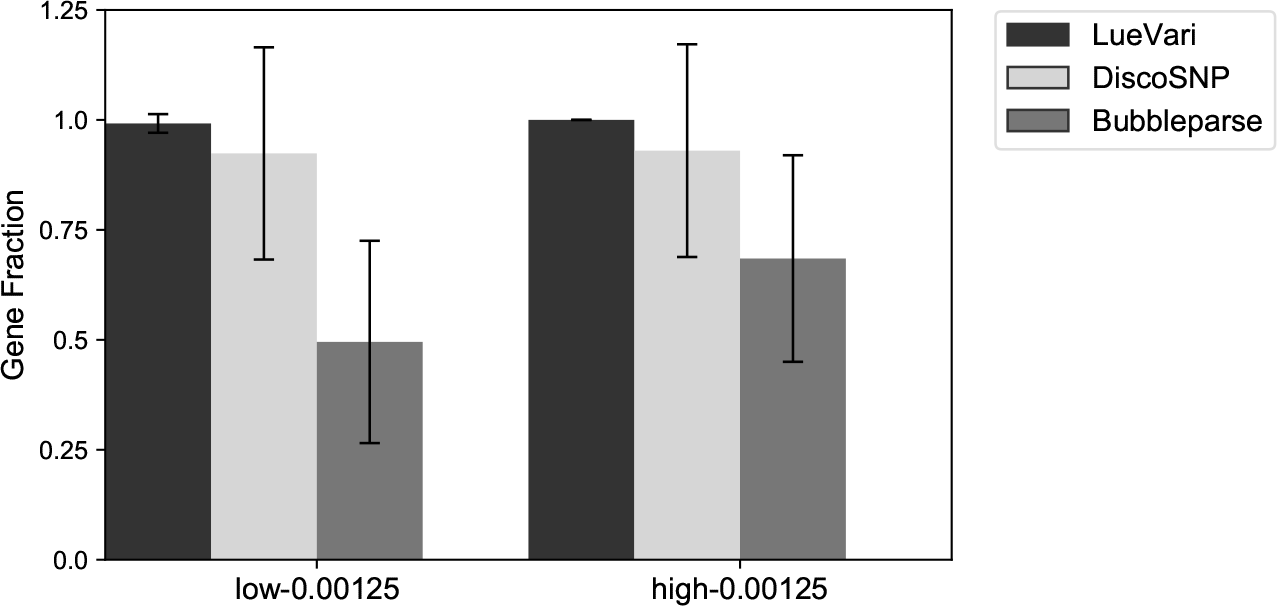
Mean gene fraction of three reference free tools on low-0.00125 and high-0.00125.

### Comparison Between Reference-Guided Methods

We used MEGARes dataset as the reference for GATK and SAMtools since they are reference-guided methods, this is an added advantage to the other methods that do not have a reference. Tables A.4 and A.5

**Fig A.4.**
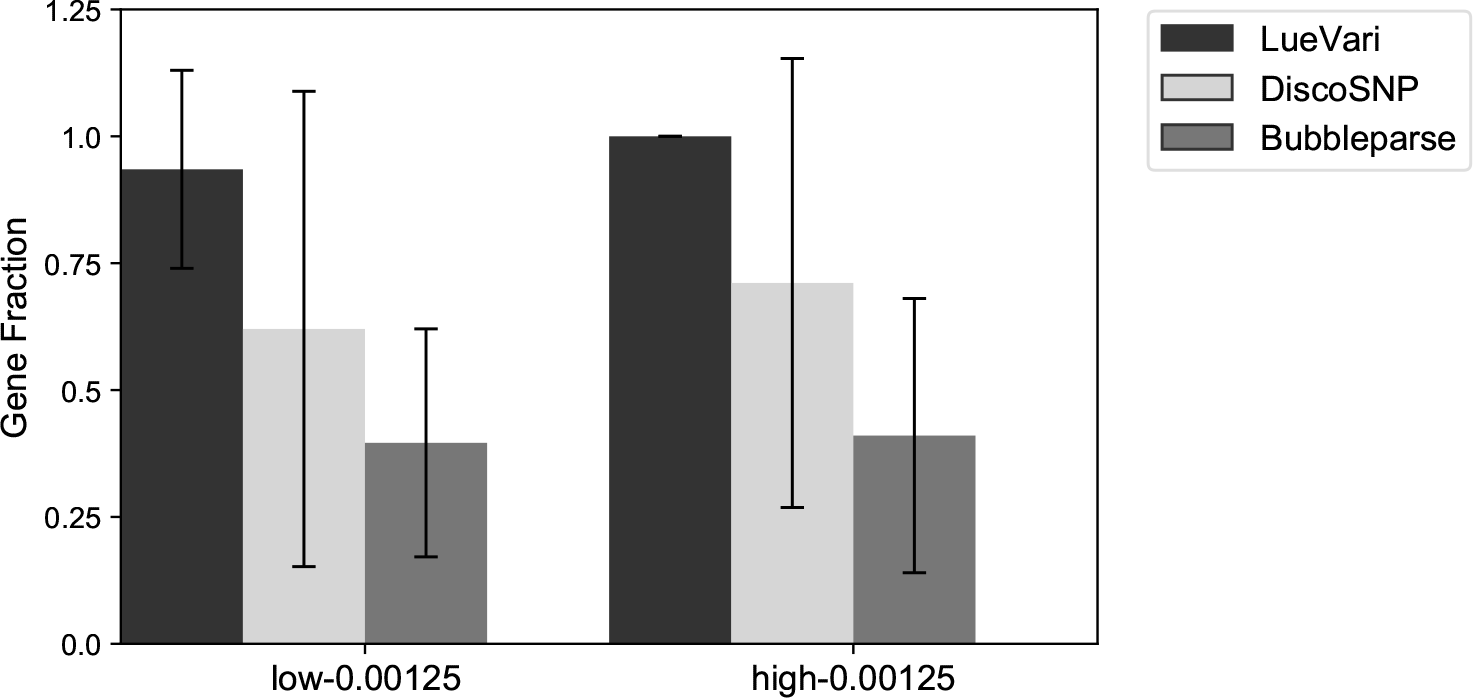
Mean gene fraction of three reference free tools on low-0.005 and high-0.005.

**Table A.4.**
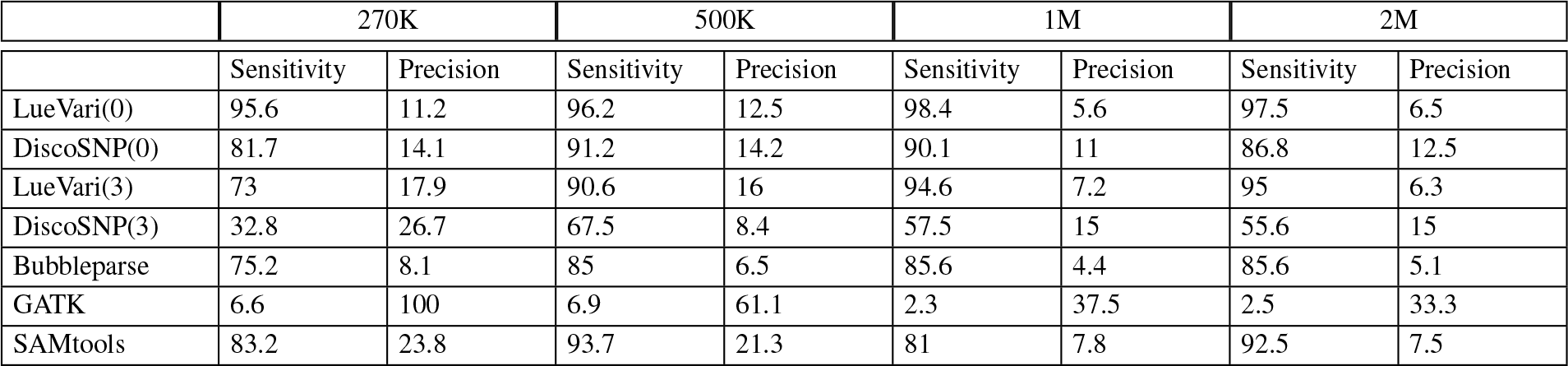
We show the accuracy of LueVari, reference-free SNP callers, and reference-guided SNP callers. We note the 270K, 500K, 1M and 2M datasets have 270,598, 527,913, 1,110,150 and 2,110,753 paired-end simulated reads, respectively. We report the sensitivity and precision as percentage. *k*=32 for all experiments except for Bubbleparse in which k=31 (k can not be even)

**Table A.5.**
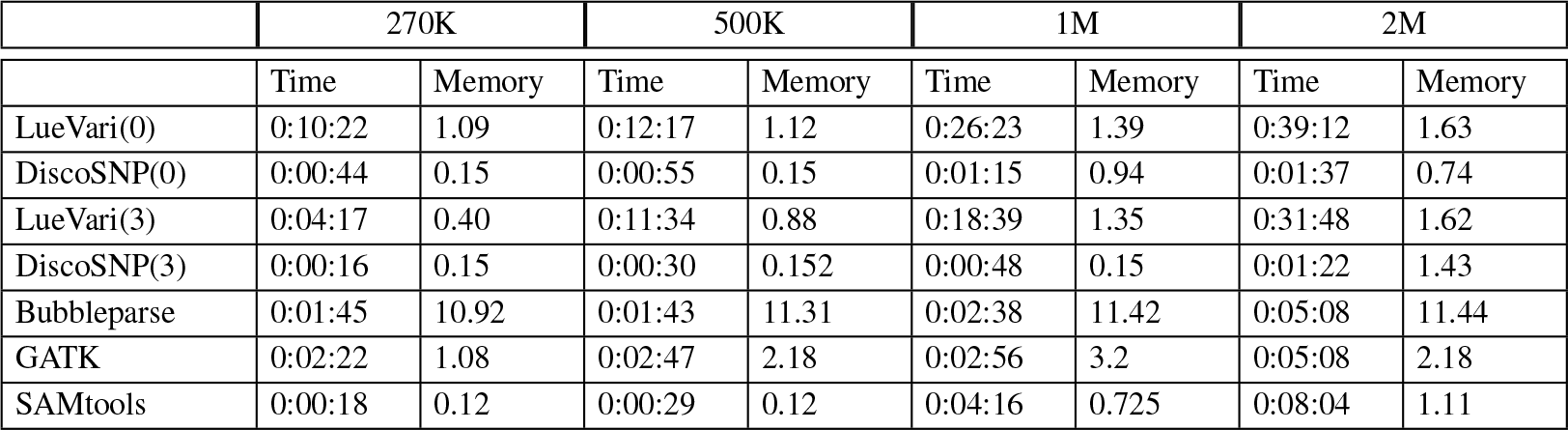
We illustrate the comparison between all competing methods. 270K, 500K, 1M and 2M with 270,598, 527,913, 1,110,150 and 2,110,753 paired-end simulated reads. We report the peak memory in gigabytes (GB), and the running time as hh:mm:ss. *k*=32 for all experiments except for Bubbleparse in which k=31 (k can not be even)

|  | 270K |  | 500K |  | 1M |  | 2M |  |
| --- | --- | --- | --- | --- | --- | --- | --- | --- |
|  | Time | Memory | Time | Memory | Time | Memory | Time | Memory |
| LueVari(0) | 0:10:22 | 1.09 | 0:12:17 | 1.12 | 0:26:23 | 1.39 | 0:39:12 | 1.63 |
| DiscoSNP(0) | 0:00:44 | 0.15 | 0:00:55 | 0.15 | 0:01:15 | 0.94 | 0:01:37 | 0.74 |
| LueVari(3) | 0:04:17 | 0.40 | 0:11:34 | 0.88 | 0:18:39 | 1.35 | 0:31:48 | 1.62 |
| DiscoSNP(3) | 0:00:16 | 0.15 | 0:00:30 | 0.152 | 0:00:48 | 0.15 | 0:01:22 | 1.43 |
| Bubbleparse | 0:01:45 | 10.92 | 0:01:43 | 11.31 | 0:02:38 | 11.42 | 0:05:08 | 11.44 |
| GATK | 0:02:22 | 1.08 | 0:02:47 | 2.18 | 0:02:56 | 3.2 | 0:05:08 | 2.18 |
| SAMtools | 0:00:18 | 0.12 | 0:00:29 | 0.12 | 0:04:16 | 0.725 | 0:08:04 | 1.11 |

### A.1.5 Results of LueVari on real datasets selected across a beef production system

We ran LueVari to identify SNPs in AMR genes in 34 samples taken from a food production facility that had previously-identified AMR genes. All experiments were performed on a 2 Intel(R) Xeon(R) CPU E5-2650 v2 2.60 GHz server with 1 TB of RAM, and both resident set size and user process time were reported by the operating system.

**Table A.1.6.**
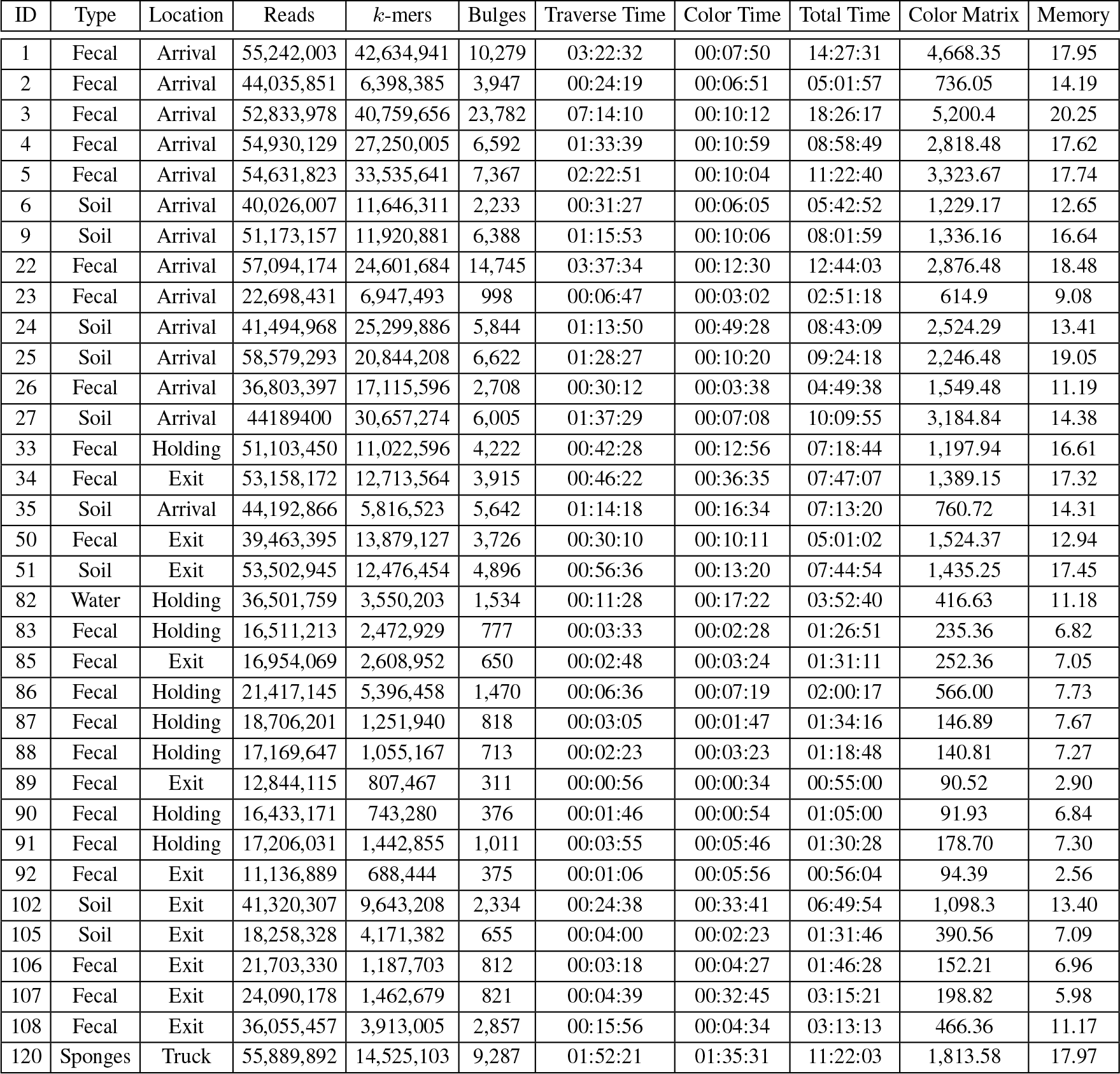
Performance of LueVari on real data samples of Noyes et al. [38]. ID, Type and the Location that each sample collected from is mentioned in first three columns. Number of Reads in each sample, number of distinct *k*-mers with frequency greater than or equal to 12, and number of found Bulges are mentioned in columns four to six. Time for Traversing the graph, time for construction of Color Matrix and Total Time are reported in columns seven to nine in format of hh:mm:ss. Size of the Color Matrix on disc is indicated in tenth column (MB), and on last column peak Memory for the whole pipeline is reported (GB).*k*=32 for all experiments except for Bubbleparse in which k=31 (k can not be even)

### Comparison of sensitivity and precision on simulated datasets with same number of reads and different copy number of the genes

In this set of experiments we simulated three datasets (the same way as what was explained in section 5.1.1), each with 300,000 paired-end reads and varying copy number of the genes. The copy numbers of the genes in these datasets are two, five and ten. We show them in table as 2, 5 and 10 respectively. As one can see in table A.7, on all experiments, LueVari has the highest sensitivity ranging from 94% to 100%.

**Table A.1.7.**
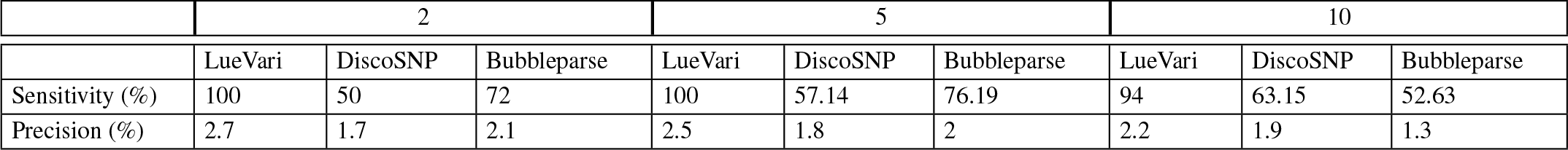
We illustrate the comparison between competing methods on datasets 2, 5 and 10 each with 300,000 paired end reads and the gene copy number of 2, 5 and 10 respectively. *k* = 32 for all experiments except for Bubbleparse in which k=31 (k can not be even). In all experiments all tools were run with the setting that achieve the highest performance (the threshold for filtering weak *k*-mers is 0).

## Footnotes

1 Reads were trimmed and those with ambiguous base calls removed

2 This is performed in an analogous way as described to find the out-degree; however, in this case we use *B_F_* rather than *B_L_*.

